## supplementary figures for "The V-ATPase-ATG16L1 axis recruits LRRK2 to facilitate lysosomal stress responses"

Figure S1

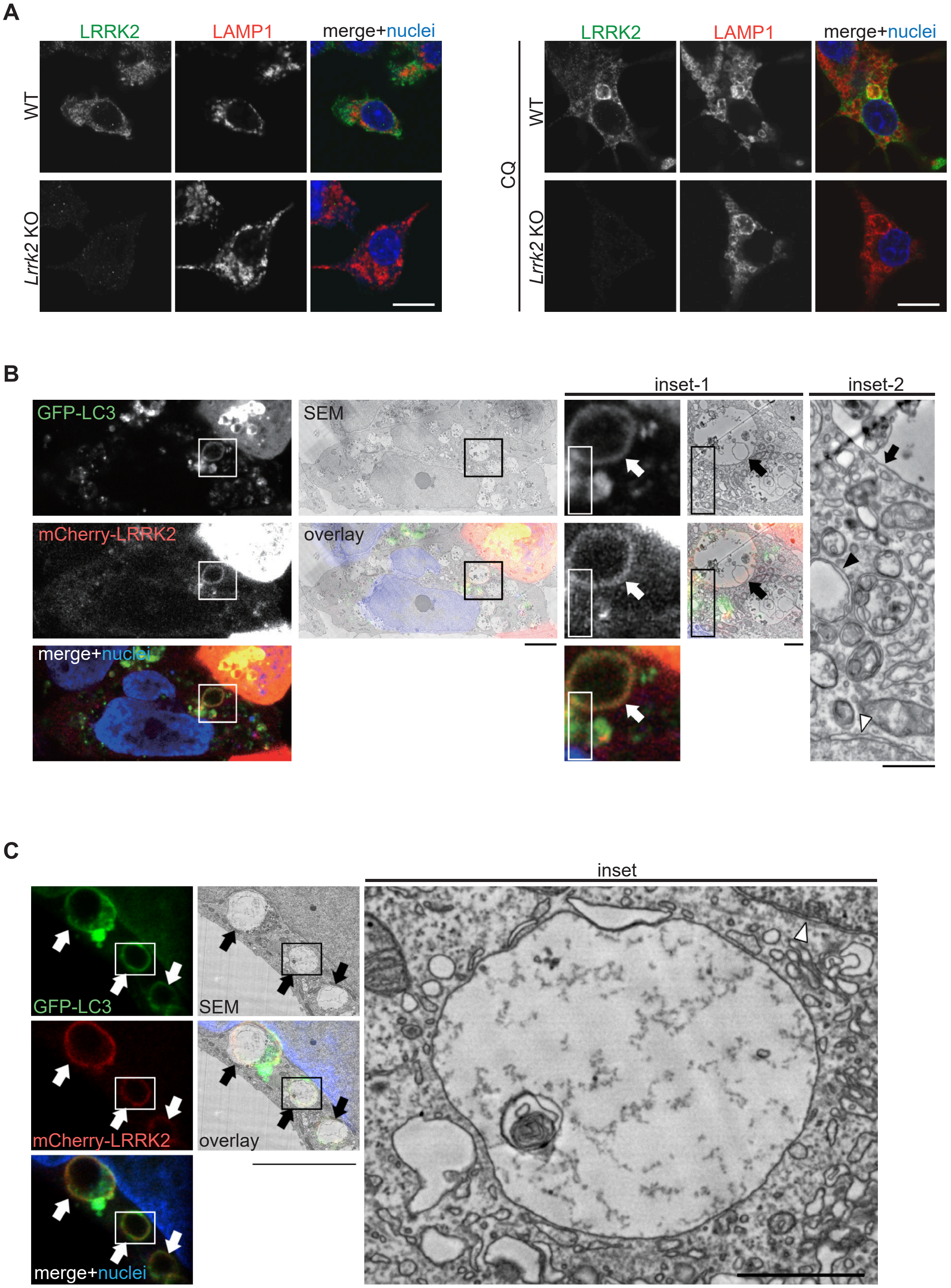


Figure S1. **Detailed analyses on LRRK2 localization. (A)** Confirmation of specificity of LRRK2 immunostaining. Immunofluorescence signal of LRRK2 stained with MJFF2 antibody was not detected in Lrrk2 knockout RAW264.7 cells in the presence or absence of CQ treatment. Scale bar, 10 μm. **(B, C)** Two representative CLEM images, in addition to Figure 1F, showing colocalization of GFP-LC3 and mCherry-LRRK2 on lysosomal single membranes under CQ treatment. Scale bars, 5 μm (B), 10 μm (C) 1 μm (inset-1 in B and inset in C), 500 nm (inset-2 in B. Arrows: LRRK2-LC3 double-positive membranes, black arrowhead in B: autolysosome, white arrowheads: nuclear envelope.

### Figure S2

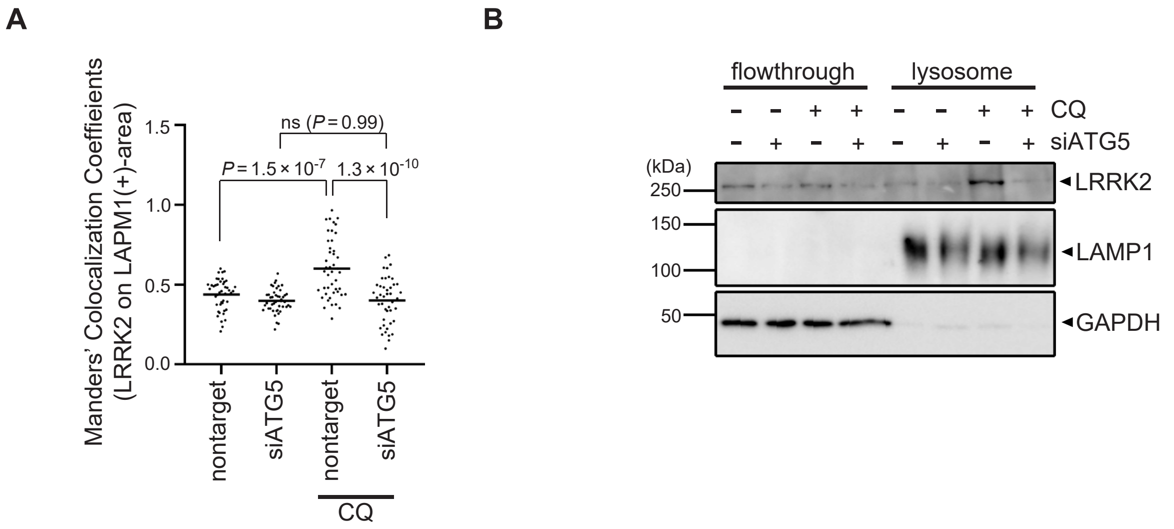


Figure S2. **Confirmation of ATG5-dependent recruitment of LRRK2 to lysosomes upon CQ treatment. (A)** Manders' Colocalization Coefficient (MCC) analysis showing the ratio of LRRK2 on LAMP1-positive area in CQ-treated and untreated RAW264.7 cells. The difference was analyzed using one-way ANOVA with Tukey's test. **(B)** Biochemical isolation of lysosomes from RAW264.7 cells showing the enrichment of LRRK2 in lysosomal fraction upon treatment with CQ but not with siATG5.

Figure S3

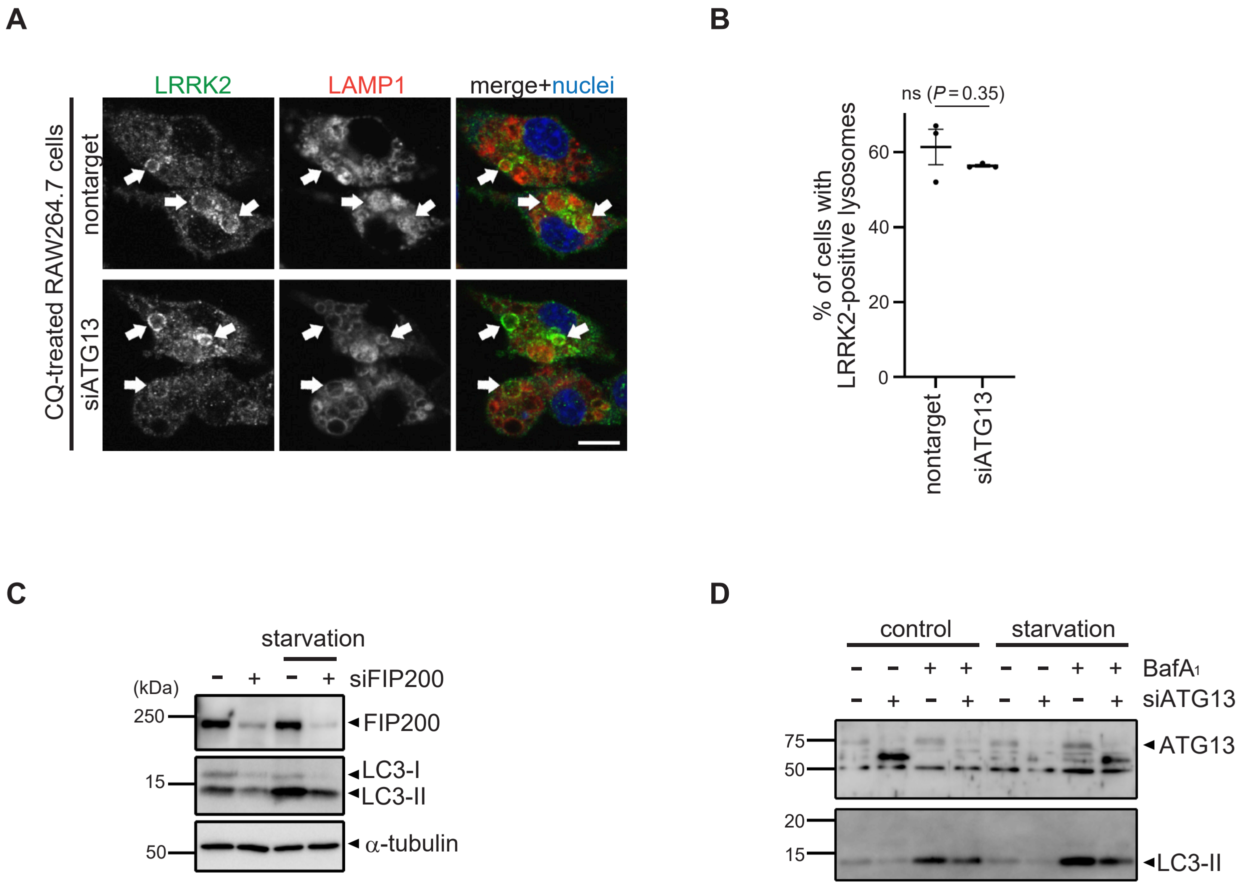


Figure S3. **Analyses of knockdown effects of ATG13 and FIP200. (A)** Fluorescence images of endogenous LRRK2 in RAW264.7 cells transfected with nontarget or ATG13 siRNA and treated with CQ. Scale bar, 10 μm. **(B)** Percentages of cells harboring LRRK2-positive lysosomes as shown in A. Data represent mean ± SEM (N = 3 independent experiments). The difference was analyzed using one-way ANOVA with Dunnet's test. ns, not significant. **(C, D)** Immunoblot pictures showing efficient knockdown of FIP200 (C) or ATG13 (D) as well as resultant decrease of starvation-induced autophagy as assessed by LC3 lipidation in RAW264.7 cells treated with siFIP200 or siATG13.

Figure S4

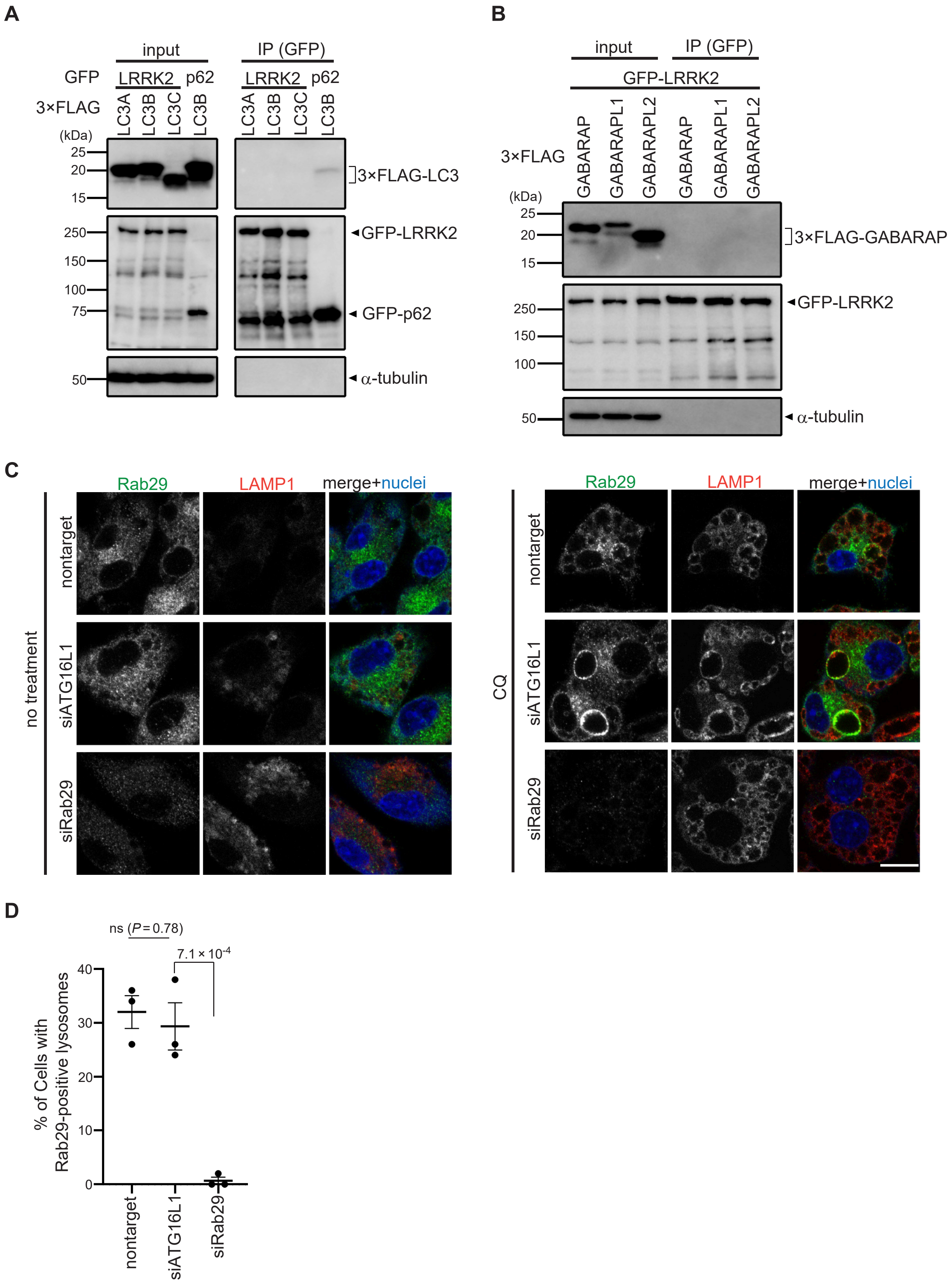


Figure S4. **Analyses on the possibility of LRRK2 recruitment via ATG8 or Rab29. (A, B)** Lack of binding between LRRK2 and ATG8 family proteins. Immunoprecipitation (IP) analysis of HEK293 cells overexpressing 3×FLAG-taggted LC3A/B/C (A) or GABARAP/GABARAPL1/GABARAPL2 (B) together with GFP-tagged LRRK2 or p62. Pulldown using GFP-trap resulted in the precipitation of LC3 with p62 but not with LRRK2. **(C)** Fluorescence images of endogenous Rab29 in RAW264.7 cells transfected with siATG16L1 or siRab29 (positive control) and treated with (right) or without (left) CQ. Arrows indicate Rab29-positive lysosomes. Scale bar, 10 μm. **(D)** Percentage of cells harboring Rab29-positive lysosomes, as shown in C. Data represent mean ± SEM (N = 3 independent experiments). The difference was analyzed using one-way ANOVA with Dunnet's test.

### Figure S5

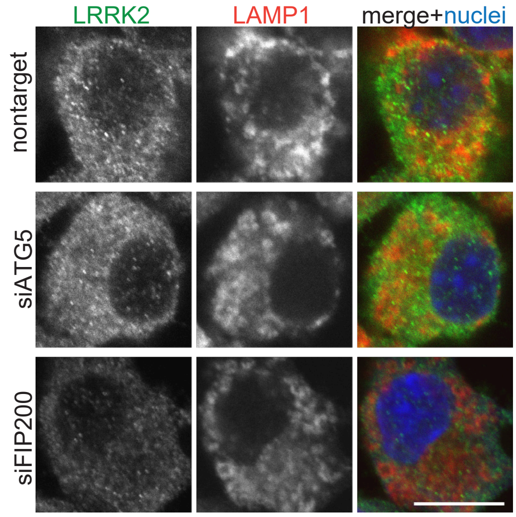


Figure S5. **Morphological analysis of lysosomes in the absence of CQ.** Represetnative fluorescence images of LAMP1-positive lysosomes as well as LRRK2 in RAW264.7 cells transfected with the indicated siRNAs, without following CQ treatment. Scale bar, 10 μm.
